# Inferring single-cell resolution spatial gene expression via fusing spot-based spatial transcriptomics, location and histology using GCN

**DOI:** 10.1101/2024.10.27.620535

**Authors:** Shuailin Xue, Fangfang Zhu, Jinyu Chen, Wenwen Min

## Abstract

Spatial transcriptomics technology allows for the detection of cellular transcriptome information while preserving the spatial location of cells. This capability enables researchers to better understand the cellular heterogeneity, spatial organization and functional interactions in complex biological systems. However, current technological methods are limited by low resolution, which reduces the accuracy of gene expression levels. Here, we propose scstGCN, a multimodal information fusion method based on Vision Transformer (ViT) and Graph Convolutional Network (GCN) that integrates histological images, spot-based spatial transcriptomics data and spatial location information to infer super-resolution gene expression profiles at single-cell level. We evaluated the accuracy of the super-resolution gene expression profiles generated on diverse tissue ST datasets with disease and healthy by scstGCN along with their performance in identifying spatial patterns, conducting functional enrichment analysis, and tissue annotation. The results show that scstGCN can predict super-resolution gene expression accurately, aid researchers in discovering biologically meaningful differentially expressed genes and pathways. Additionally, scstGCN can segment and annotate tissues at a finer granularity, with results demonstrating strong consistency with coarse manual annotations.

**Key Points:**

- scstGCN combines multi-modal information including histology image, spot-based spatial transcriptomics (ST) data, and physical spatial location through deep learning methods to achieve single-cell resolution of spot-based ST data without requiring single-cell references.
- scstGCN employs GCN to capture complex relationships between neighboring cells, facilitating the integration of multimodal feature information based on single-cell level, and then accurately infers single-cell resolution spatial gene expression.
- scstGCN can infer single-cell resolution gene expression across the entire tissue region. Through transfer learning, gene expression in three-dimensional tissues can be characterized efficiently. Furthermore, it demonstrates outstanding performance in spatial patterns enhancement, functional enrichment analysis, and annotate tissues at the high-resolution.

## Introduction

In biology, the spatiotemporal specificity of gene expression dictates that integrating cellular spatial location information is essential for a better understanding of the specific functions of cells within tissues [1]. Neglecting the spatial information of cells may lead to inadequate understanding of gene expression patterns and spatial distribution. However, the spatial location information of tissue cells cannot be retained in single-cell sequencing experiments due to technical limitations [2, 3]. With the innovation of spatial transcriptomics (ST) technology, it has become possible to detect the quantity of gene transcripts within tissues while retaining spatial location information [4]. Many ST techniques have been developed to reveal the spatial heterogeneity of cells and genes and their distribution patterns in tissue space [5]. Therefore, ST technologies have been used in research on the development of tissues and organs [6], neurology [7] and tumor heterogeneity [8].

At present, the main ST techniques can be divided into two categories: (1) imaging-based techniques, including in situ hybridization and in situ sequencing [9]. In situ sequencing-based approach is to reverse-transcribe RNA in a cell by targeting probes, followed by sequencing using a DNA ligase-based sequencing approach, such as FISSEQ [9] and STARmap [10]. In situ hybridization is another imaging-based method that uses fluorescently labeled nucleic acid probes to determine the spatial location and abundance of DNA and RNA in tissues and cells, typically MerFISH [11, 12] and seqFISH [13]. (2) next-generation sequencing (NGS)-based approaches, works by fixing oligonucleotide sequences carrying spatial barcodes to a chip at a specific resolution, tissue sections were then affixed to the chip for cell lysis and RNA construction Library sequencing [14], such as ST and 10X Genomics Visium. However, each of these technologies has its drawbacks. Imaging-based technologies can provide higher resolution and sensitivity, but have lower throughput, a large number of transcriptomes can not be detected. In principle, NGS-based technologies can detect a complete tissue atlas and capture all transcriptome information, but it is limited by lower spatial resolution to more accurately study detailed gene expression patterns. 10X Visium is currently the most widely used ST technology. However, on the 10X Visium platform, the center-to-center distance between spots is 100 *µ*m and the spots with a diameter of 55 *µ*m may contain 5 to 30 cells. Even on the older ST platform, which has 100 *µ*m spot diameter with 200 *µ*m center-to-center distance, the number of cells within a spot may up to 200. Therefore, although the availability of many ST platforms, none of them provide a comprehensive solution that balances resolution and sequencing depth.

Recently, some tools have been developed to address the various shortcomings of ST technology using deep learning-based methods. For example, some spatial clustering methods have been developed to help identify spatial domains, such as AVGN [15], STAGATE [16], and STMask [17], using graph neural networks as the main framework to accurately capture spatial domain information. In addition, a few ST data imputation techniques, like SpatialScope [18] and SpaDiT [19], have been designed to improve the low sensitivity of ST data. Furthermore, spot-level gene expression prediction methods, such as BLEEP [20] and mclSTExp [21], aim to predict spot-based spatial gene expression from histology images to eliminate the cost-prohibitive of ST field.

In ST technology studies, enhancing the resolution of spatial gene expression is a core task. Low spatial resolution and incomplete tissue coverage may hinder researchers from identifying gene expression patterns and deeply characterizing intricate tissue architectures. Ideal ST data should have single-cell resolution and cover the entire tissue surface. Enhancing the resolution of spot-based ST data to the cellular level can effectively solve the above problems and help discover more biologically significant pathways.

Several methods have been developed to enhance the resolution of spatial gene expression by imputing the gaps between spots, dividing spots into sub-spots, or mapping the entire tissue into a collection of superpixels representing single cells. For example, stEnTrans [22] establishes self-supervised tasks to predict gene expression in gaps between spots using the Transformer encoder. STAGE [23] utilizes a spatial location-supervised auto-encoder to predict high-density gene expression. BayesSpace [24] uses Bayesian modeling to estimate gene expression data at the subspot level without relying on histological images. However, none of these methods have achieved comprehensive prediction of single-cell resolution gene expression across the entire tissue. Xfuse [25] integrates ST data and histology images using a deep generative model to infer super-resolution gene expression profiles. TESLA [26] generates high-resolution gene expression profiles based on Euclidean distance metric, which considers the similarity in physical locations and histological image features between superpixels and measured spots. However, they have not deeply extracted the intrinsic features of histological images and inefficiently utilized existing data. iStar [27] employs a multi-layer Transformer architecture to extract features from histological images, aiming to predict super-resolution gene expression by capturing both local patterns and global spatial relationships, but it does not consider for the strong correlations between neighboring cells clusters, failing to capture the complex relationships between adjacent superpixels. Additionally, gene expression correlates with spatial location, a factor overlooked by all methods predicting super-resolution gene expression.

Here, we developed scstGCN to robustly predict super-resolution closed to single-cell spatial gene expression with the ViT module and the multimodal-features-supervised GCN module by integrating histological image features, spatial positional information, and spot-based ST data. Compared to existing inferring super-resolution methods, scstGCN integrates histological image features and spatial location information, leveraging GCN to effectively capture complex relationships among adjacent superpixels. Comprehensive benchmarking on ST data generated on multiple diverse disease and healthy tissues from different platforms demonstrates the exceptional super-resolution gene expression prediction capability of scstGCN. Moreover, scstGCN has exhibited its capability in enhancing spatial patterns of significant genes, discovering more biologically meaningful pathways and high-resolution annotation of tissue architecture.

## Materials and methods

### Dataset description and preprocessing

To precisely evaluate the performance of scstGCN, we used multiple Xenium datasets including human breast cancer (HBC), human pancreas (HP), and hHeart Non-diseased (HN) generated by 10x Genomics. Xenium dataset contains subcellular-based ST data, along with spatial positional information of the subcellular locations. We preprocessed Xenium datasets to align with our experimental objectives. To obtain ground truth, we constructed a rectangle grid of superpixels and then partitioned the cell-level gene expression into this rectangle grid based on the overlap area between cells and the grids:

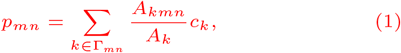

where *p*_*mn*_ and *c*_*k*_ represent the gene expression at superpixel *P*_*mn*_ and cell *C*_*k*_, respectively; *A*_*k*_ and *A*_*kmn*_ represent area of cell *C*_*k*_ and its overlapping area with the superpixel *P*_*mn*_, respectively; Γ_*mn*_ represents the set of all cells that overlap with the superpixel *P*_(_*m, n*). Next, pseudo-Visium data was obtained by segmenting the gene expression of Xenium dataset into a series of spots based on the spot size and center-to-center distance of the Visium:

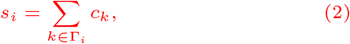

where *s*_*i*_ represents the gene expression of pseudo-spot *S*_*i*_, and Γ_*i*_ represents the set of all cells covered by pseudo-spot *S*_*i*_. The size of each superpixel was set to 8*×*8 *µ*m^2^, which is approximately equivalent to the size of a single-cell. The spot diameter was set to 55 *µ*m, and the center-to-center distance was set to 100 *µ*m. The location information of the pseudo-spots are evenly covered on the entire detection tissue based on the characteristics of Visium data.

Visium HD technology is 10X Genomics’ latest high-resolution spatial transcriptome high-throughput sequencing technology. It enhances the resolution to the single-cell level while maintaining the advantages of high gene throughput. Unlike Xenium data, it can detect more genes, and the sequencing unit is a square area without gaps (such as 8*×*8 *µ*m). Therefore, processing Visium HD data only requires simulating pseudo-Visium data. We need to segment the super-resolution gene expression of Visium HD data into a series of spots based on the spot size and center-to-center distance of the Visium to obtain the pseudo-Visium data. In simple terms, the gene expression of the sequencing unit set covered by the spot is aggregated as the gene expression of the spot:

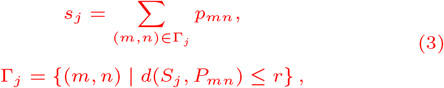

where *p*_*mn*_ and *s*_*j*_ represent the gene expression at sequencing unit *P*_*mn*_ and pseudo-spot *S*_*j*_, respectively; *d*(*s*_*j*_, *p*_*mn*_) represent the center distance between pseudo-spot *S*_*j*_ and sequencing unit *P*_*mn*_; *r* represent the radius of the Visium data. The location information of the pseudo-spots are evenly covered on the entire detection tissue based on the characteristics of Visium data.

Additionally, we also evaluated scstGCN numerically on the human dorsolateral prefrontal cortex tissue (DLPFC) conducted from 10X Visium platform. Since the resolution of DLPFC data is at the spot-level, and the resolution of our predicted gene expression is close to the cell-level, we merged the predicted single-cell resolution gene expression into the spot-level gene expression based on the location information of all spots in the original DLPFC data. This process is basically the same as the process of generating pseudo-Visium data from Visium HD. The difference is that the location information of pseudo-spot *S*_*j*_ comes directly from the original data, spot 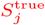. Flow of preprocessing for all single-cell resolution datasets can be inferred in Supplementary Figure S1.

For all Xenium datasets, we predicted all genes in each dataset (313 genes in HBC S1R1 and HBC S1R2, 288 genes in HBC S2, 377 genes in HP and HN). For all spot-based ST datasets and Visium HD datasets, we selected the top 1000 highly variable genes as experimental samples. Other details of description for all datasets can be found in Table 1 and Supplementary Table S1.

**Table 1.** Summary of ST datasets used in this study. Among them, MBHD, HBCHD, HP, HN, MBC, and MBS datasets were also collected from the 10x Genomics portal.

| Datasets | Size/Radius | Sections | Genes | Platform |
| --- | --- | --- | --- | --- |
| MBHD | single-cell | 1 | 17,797 | Visium HD |
| HBCHD | single-cell | 1 | 18,085 | Visium HD |
| HP | single-cell | 1 | 377 | Xenium |
| HN | single-cell | 1 | 377 | Xenium |
| MBC | 55um | 1 | 19,465 | 10X Visium |
| MBS | 55um | 1 | 32,285 | 10X Visium |
| HBC [28] | single-cell | 3 | 288-313 | Xenium |
| HER2ST [29] | 100um | 32 | 10,530 | ST |
| DLPFC [30] | 55um | 12 | 33,538 | 10X Visium |

### Overview of scstGCN

scstGCN is specifically designed to infer super-resolution gene expression by integrating histological image features, spatial position information, and spot-based ST data. We first extract multimodal feature map, and then use the GCN module to predict single-cell resolution gene expression from multimodal feature map by a weakly supervised framework (Figure 1).

**Fig. 1.**
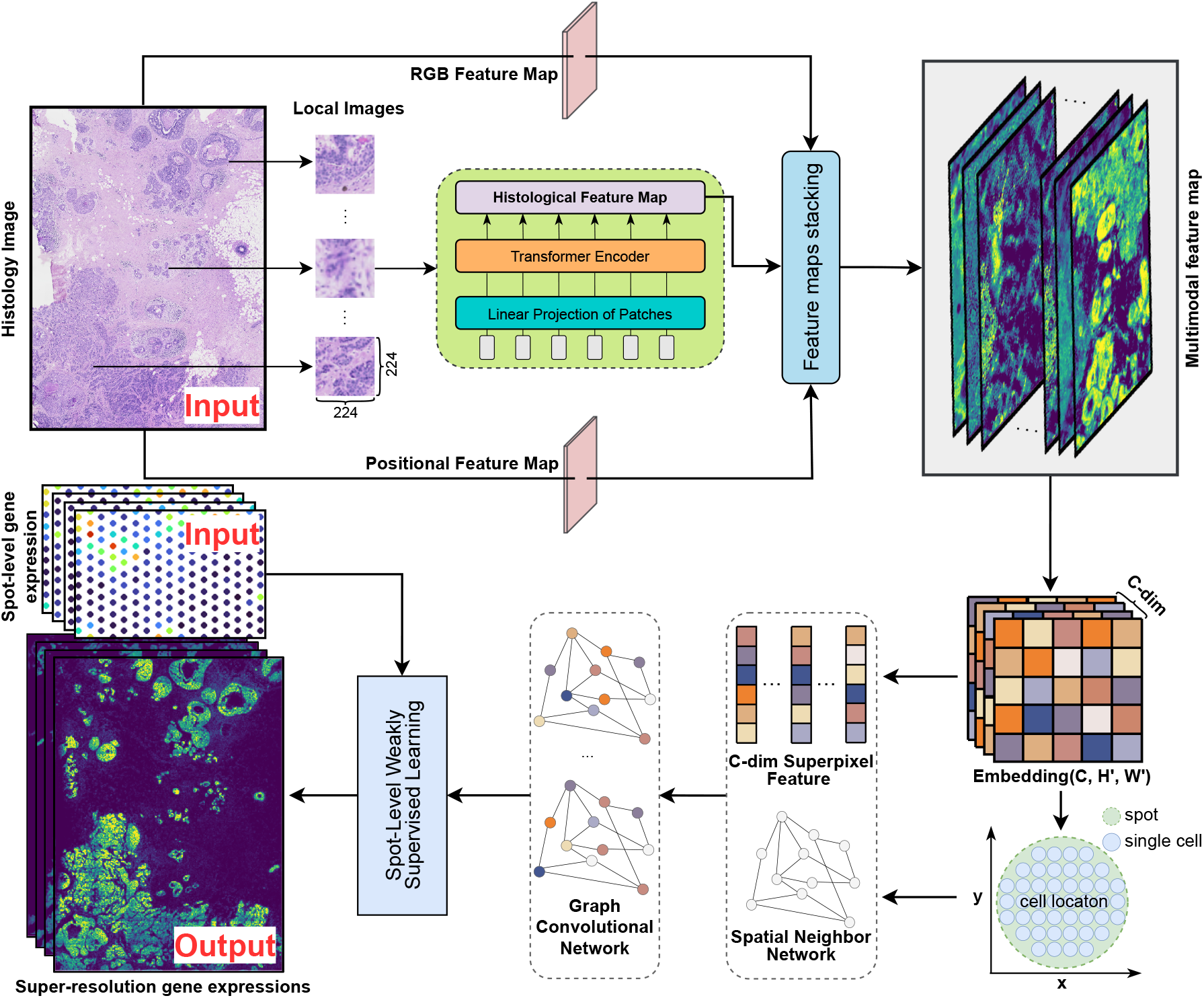
The architecture summary of scstGCN. First, the histological image is divided into sub-images of size 224 to fit the input of the ViT module, which can extract histological feature maps by converting image data into sequence data. Second, the RGB feature map can be obtained from the histological image through downsampling and then the location feature map is calculated based on the two-dimensional spatial coordinates of the superpixels. These features are stacked to obtain the multimodal feature map. Next, The GCN module is utilized to further capture the complex relationships between adjacent cells. Finally, based on a weakly supervised GCN framework, the original gene expression data is employed as pseudo-labels to predict super-resolution gene expression.

To obtain histological feature map, scstGCN first divides the histological image into a series of subimages, and then extracts histological image features using the ViT module. The positional feature map is constructed based on the two-dimensional coordinates of the superpixels, while the Red-Green-Blue (RGB) feature map is obtained through simple downsampling. Next, multimodal feature map is obtained by stacking the histological feature map, positional feature map, and RGB feature map (Figure 1). Generally, gene expression is largely influenced by complex interactions between adjacent cells. To model this, scstGCN adaptively represents the intricate correlations between neighboring superpixels using the GCN modules, thereby effectively establishing the relationship between multimodal feature map and super-resolution gene expression (Figure 1). We adopt a weakly supervised learning framework to train scstGCN because the model outputs are at the superpixel-level while the training data are at the spot-level. Specifically, we model the sum of gene expression of the superpixels covered by spots and the gene expression of spots.

### Extracting multimodal feature map based ViT module

A substantial body of study has confirmed the strong correlation between histology images and gene expression patterns [31, 32]. ViT has demonstrated remarkable success in image analysis. Therefore, we aim to leverage ViT to extract comprehensive features from histology images. Due to varying pixel sizes in histology images from different datasets, we need to resize the histology images, defaulting to scaling the pixel size to 0.5 *µ*m. This process ensures that 16*×*16 patches approximately correspond to the size of a single cell. When the scaled image is not divisible by 224, padding operation is required for the image so that its height and width are both divisible by 224.

Next, we partition the entire histological image into subimages of size 224*×*224 to serve as inputs to the ViT module to extract histology image features in tiles. Let *M* and *N* denote the height and width of the histology image, then it can be expressed as 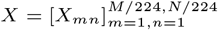, where *X* ∈ ℝ^*M*^ *×* ℝ^*N*^ *×* ℝ^3^ and each subimage *X*_*mn*_ ∈ ℝ ^224^ *×* ℝ ^224^ *×* ℝ ^3^.

Following the traditional principles of ViT [33], we reshape each subimage into a series of 16*×*16 patches, and then flatten the image into sequence data for compatibility with the Transformer architecture:

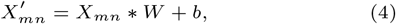

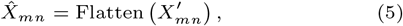

where 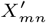 and 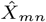 have shapes (16*×*16*×*3,224/16,224/16) and (196, 768); *W* and *b* represent the filter parameters and biases of the convolutional structures.

Next, we briefly describe the basic components of the Transformer block, including Multi-head Self-Attention (MSA) and Multi-Layer Perceptron (MLP).

In the MSA module, the inputs *U* ∈ ℝ^*n×d*^ are linearly transformed into three parts, namely 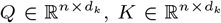, and 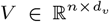 where *n* is the sequence length, *d, d*_*k*_, and *d*_*v*_ are the dimensions of inputs, keys (queries) and values respectively. Linear transformation can be described as follows:

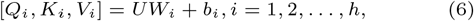

where *h* denotes the numbers of self-attention operations in MSA, *W*_*i*_ and *b*_*i*_ represent parameters and biases of the linear transformation. The scaled dot-product attention is applied on *Q, K, V* and then concat all head as the output:

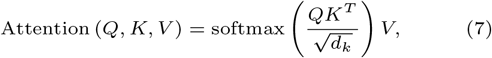

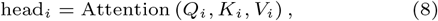

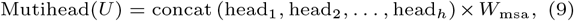

where 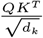 represents the similarity between Query and *W*_msa_ is the weight matrix for multiple heads.

There are two linear transformations separated by an activation function in the MLP module for feature transformation and non-linearity:

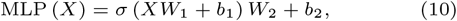

where *W*_1_, *b*_1_, *W*_2_, and *b*_2_ are the weight and bias term of the first and second fully-connected layer respectively; *σ*represents the activation function such as GELU [34].

We use the function *f* (•) to represent the above process of feature extraction using the ViT module. ViT maps each 16*×*16 patch within each subimage *X*_*mn*_ into a feature vector of length 1024, and then the histological image features are obtained by applying the inverse operation of Flatten:

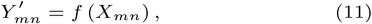

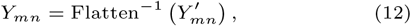

where 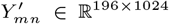 sequential data, and *Y*_*mn*_ ∈ ℝ^1024*×*14*×*14^ is 3-D image data. After extracting features from all subimages, the entire histological feature map 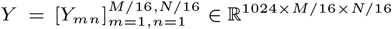 are obtained.

Additionally, certain regions within the tissue may exhibit similar histological features while differing in gene expression levels. By incorporating spatial location features, we aim to enhance the robustness and accuracy of the model. We calculate the positional feature map *P* ∈ ℝ^2*×M/*16*×N/*16^ based on the two-dimensional spatial location information of the superpixels:

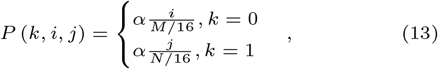

where *i* and *j* represent the two-dimensional coordinates of superpixel, and the hyperparameter *α* is set to be 1 in the experiments.

Finally, we obtain the RGB feature map *T* ∈ ℝ^3*×M/*16*×N/*16^ by resizing the entire histological image to the desired size *M/*16 *× N/*16 using average pooling. From this point, we have obtained histological feature map *Y*, positional feature map *P*, and RGB feature map *T*. By stacking these feature maps along the channel dimension, we obtain the multimodal feature map:

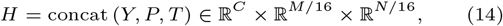

where *H* can be represented as 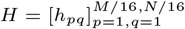, with each superpixel *h*_*pq*_ ∈ ℝ^*C*^ is the feature vector at pixel (*p, q*). *C* = 1024 + 2 + 3, which is the sum of the channels of the feature maps *Y, P*, and *T*.

Additionally, due to the benefits of transfer learning in ViT architecture, we pretrain ViT module through self-supervised learning (SSL). This step only requires histological images, making many publicly available histopathology datasets suitable for model pretraining. In our experiments, we utilized the general pathology self-supervised model named UNI [35], which uses DINOv2 [36] to pretrain on over 100 million images from diagnostic H&E-stained WSIs spanning 20 major tissue types.

### Inferring super-resolution gene expression using GCN module

Our purpose in extracting multimodal feature map is to use it for inferring super-resolution gene expression profiles. In tissue structures, the gene expression exhibits a high degree of correlation with adjacent cells, far exceeding that with regions of tissue that are physically distant from each other. GCN module is well-suited for establishing complex communication relationships between neighboring cells. Therefore, we capture complex relational information between adjacent superpixels using an undirected graph *G*(*V, E*) when predicting gene expression at the single-cell level. Each vertex *V* symbolizes the superpixel, characterized by a multimodal feature vector *h*_*pq*_. And the edge *E* represents the connections between two vertices. Due to the large number of superpixels, we consider the set of superpixels spanned by each spot as a graph to reduce computational complexity. Let *S* be the number of spots in the original ST data, then training data 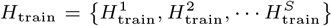 where 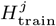 is the set of superpixels spanned by spot *i* of size *D × D* with *C* channels. Here, *D* denotes the number of superpixels covered by the spot’s diameter. Next, we define each superpixel as a node, and select the four closest nodes based on physical distance as its neighbors to generate the adjacency matrix *A*. After two layers of GCN, which fuse information between adjacent cell-level, a 512-dimensional feature vector at each superpixel is obtained. The above process can be described as follows:

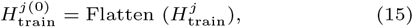

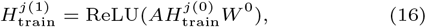

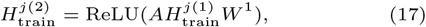

where 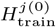 has shape 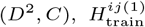 and 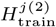have the same shape (*D*^2^, 512), with *D*^2^ represents the number of nodes for graph.

Next, 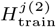 are input into the output layer containing a linear layer with 512 input nodes and *K* output nodes, where *K* represents the number of genes. The outputs are activated by an exponential linear unit (ELU) [37] to ensure that the predicted super-resolution gene expressions are non-negative. The process of transferring from feature vectors to gene expressions is as follows:

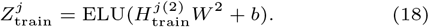

To train GCN module and output layer, due to training data lacks high-resolution gene expression profiles as labels, we employ a weak supervision framework. Specifically, we model the gene expression of each spot as the sum of the gene expressions of the superpixels covered by that spot. Regions not covered by any spot are excluded from the training process. Let 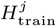 be the set of superpixels spanned by spot *j, F* (*·*) represents the network composed of the GCN module and the output layer, 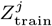 be the predicted super-resolution gene expressions corresponding to 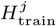, Filter(*·*) represents the operation of filtering out superpixels not contained within round area of spots, *g*_*j*_ be the observed gene expressions at spot *j*. Then the weak supervision framework can be described as:

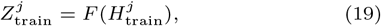

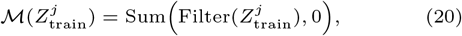

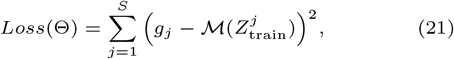

where *g*_*j*_ and 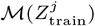 are both spot-level gene expression vectors of length *K*.

We use ADAM optimizer to minimize the loss via mini-batch gradient descent. After completing the training phase, we neatly divide the entire multimodal feature map into a series of graphs as predicting data 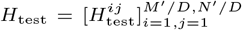 where *M*′ = *M*/16 and *N*′ = *N/*16. And then we can infer super-resolution gene expression use the trained network *F* (*·*):

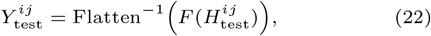

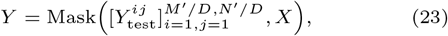

where the shape of *Y* is (*M*′, *N*′, *K*), with each *Y* ^*ij*^ has shape (*D, D, K*). Function Mask(*·*) first detects the contour of the captured tissue regions in the histological image *X*, then masks out values outside that detected contour in super-resolution gene expression data *Y*. The *k*-th channel of *Y* with shape (*M*′, *N*′) represents the super-resolution gene expression profile predicted by scstGCN for the gene *k*.

## Results

The details on the baseline methods, implementation details, and evaluation metrics can be found in Supplementary Note 1, 2 and 3.

### scstGCN can better predict super-resolution gene expression profiles

Due to the lack of single-cell resolution in spot-based ST data, it is not possible to perform precise quantitative evaluation of predicted super-resolution gene expression profile. To assess the accuracy of scstGCN in predicting super-resolution gene expression, our numerical evaluation experiments were conducted on multiple simulated datasets derived from Xenium data.

First, we applied scstGCN and other methods on pseudo-Visium dataset derived from the HBC dataset. HBC comprises two samples: Sample 1 includes two consecutive sections (denoted as HBC S1R1 and HBC S1R2), while Sample 2 contains only one section (denoted as HBC S2). We compared the prediction accuracy of scstGCN with iStar [27], XFuse [25], TESLA [26], and STAGE [23]. Some methods were excluded such as BayesSpace [24] and stEnTrans [22] because BayesSpace separates a spot into several sub-spots and cannot impute gene expression for unmeasured locations; stEnTrans merely interpolated the gaps between spots and did not enhance the gene expression levels to super-resolution. We calculated root mean square error (RMSE) and structural similarity index measure (SSIM) [38] between ground truth and each predicted super-resolution gene expression. The results show that scstGCN attains lower median of gene-wise RMSE and higher median of gene-wise SSIM across all sections in HBC (Figure 2). scstGCN’s 75th percentile (0.055, 0.056, 0.062 respectively) of RMSE is lower than the minimum of 50th percentiles of iStar (0.073, 0.063, 0.067 respectively) in HBC (Figure 2A). What’s even weirder is that the 25th percentile of SSIM for scstGCN is higher than or comparable to the maximum 75th percentile of SSIM for iStar in HBC (Figure 2B). Additionally, to demonstrate scstGCN’s broad applicability, we conducted the same experiment on HP and HN datasets (details in Table 1), and the results were consistent with those observed in HBC data, with scstGCN achieving the best performance (Figure 2).

**Fig. 2.**
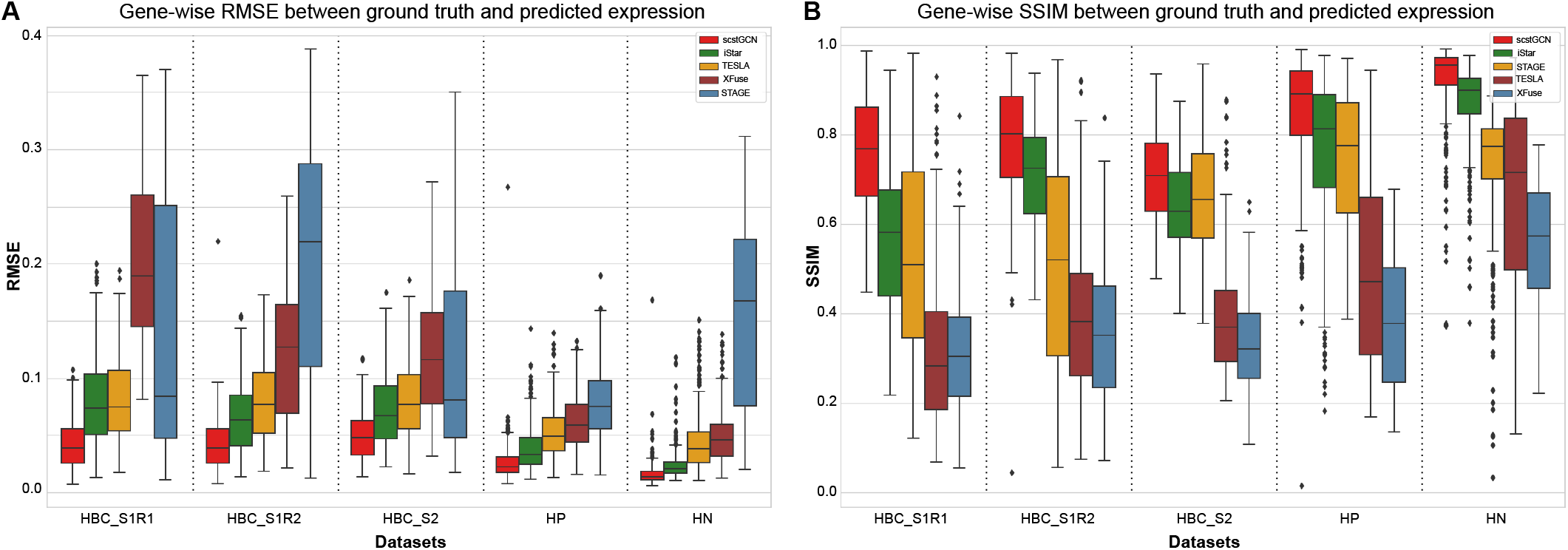
Results on multiple Xenium ST datasets demonstrate scstGCN can predict super-resolution gene expression with higher accuracy. (**A**) Calculation result in term of the Root Mean Square Error (RMSE) metric. (**B**) Calculation result in term of the Structural Similarity Index Measure (SSIM) metric. We calculated RMSE and SSIM metrics between ground truth and predicted super-resolution gene expression using scstGCN, iStar, TESLA, XFuse and STAGE on multiple ST datasets.

To intuitively analyze the predictive capabilities of different methods, we spatially visualized several genes that have different spatial patterns. Figure 3 shows the comparison of pseudo-Visium, ground truth and the predicted super-resolution gene expression using scstGCN, iStar, XFuse and TESLA about certain genes. Visually, the predictions from scstGCN are the closest to the ground truth measured by Xenium data. Despite achieving the same super-resolution as scstGCN, iStar predicts high gene expression in certain tissue areas (such as globally highly expressed in *ERBB2*, partially highly expressed in *KRT8* and *PTPRC*), XFuse exhibits distortion in genes with weak spatial patterns, and TESLA can only roughly predict gene expression patterns in all genes. Finally, to illustrate that scstGCN has powerful transfer learning capabilities, the pseudo-Visium data of HBC S1R2 were used as the training data, and super-resolution gene expression profiles was obtained on HBC S1R1 using only its histological image as the input (Supplementary Figure S2). The results show that both in terms of evaluation metrics and spatial expression analysis, scstGCN outperforms the state-of-the-art method iStar.

**Fig. 3.**
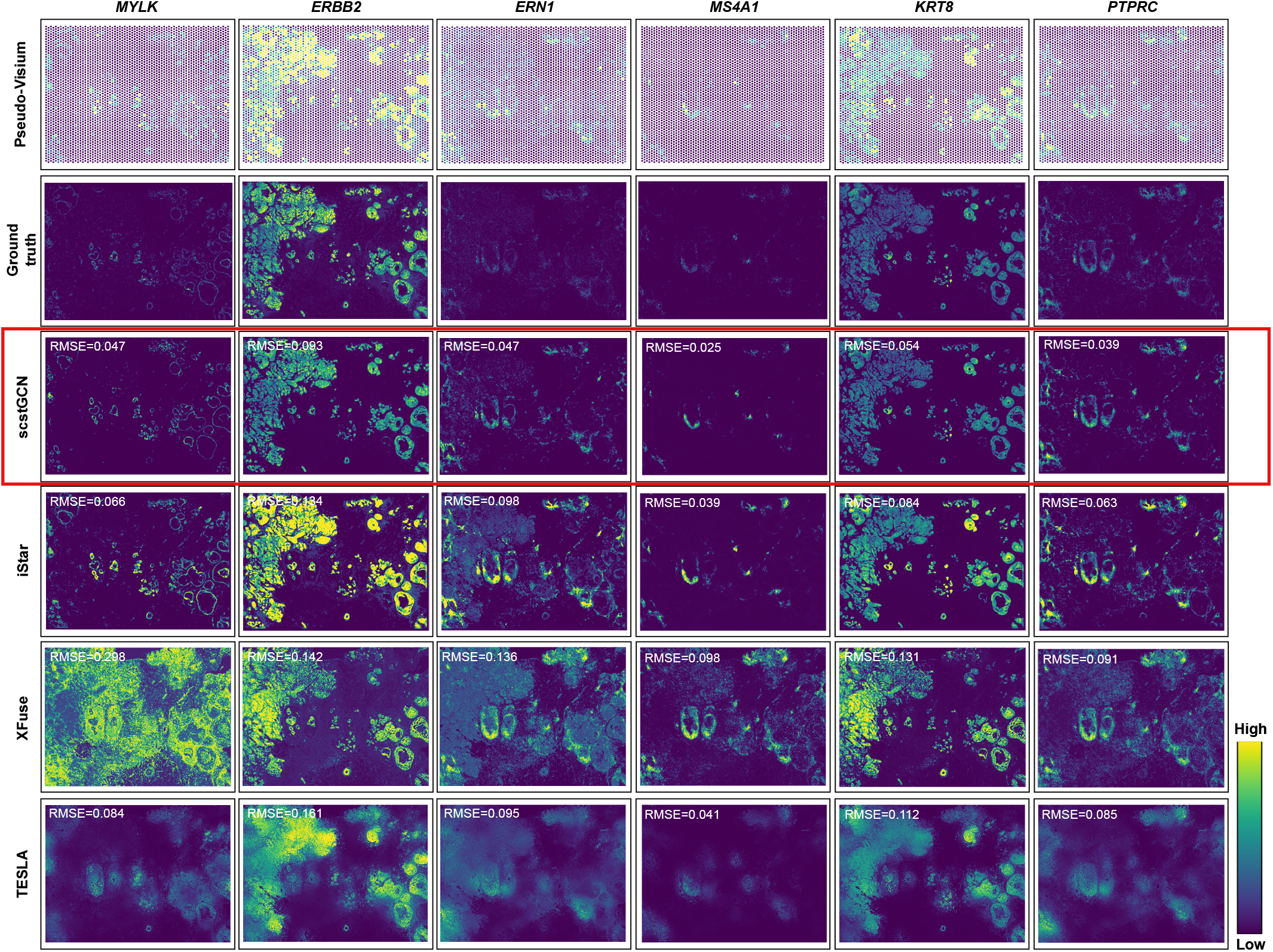
Spatial expression analysis of multiple groups of genes with different spatial patterns in HBC S1R1 data further demonstrate the superiority of scstGCN. The analysis are based on pseudo-Visium data, super-resolution ground truth, and predicted data using scstGCN, iStar, XFuse and TESLA. Each column corresponds to a gene, with the first two rows from the top displays the pseudo-Visium and ground truth, while the subsequent rows show the predicted data using different methods.

### scstGCN enables accurately restore gene expression data of Visium HD technology

Recently, 10X Genomics has launched Visium HD technology, which enhances the resolution to the single-cell level while maintaining the advantages of high gene throughput. Compared to Xenium, Visium HD has two advantages for our study: (1) the sequencing unit of Visium HD is a square area without gaps (8*×*8 *µ*m), which is consistent with the output of our task. This character eliminates the need to map scattered cellular gene expression onto a rectangle grid of superpixels, enhancing the reliability of the experimental results. (2) Visium HD offers the advantage of high gene throughput, allowing us to evaluate the performance of the model on a greater number of genes. In view of the above benefits that are more capable of verifying the accuracy of our method, we selected two Visium HD datasets: Human Breast Cancer (HBCHD) and Mouse Brain (MBHD).

We used the 8*µ*m bins of the Visium HD data as ground truth, which is approximately equal to single-cell resolution. Similar to the simulation process on Xenium data, pseudo-Visium data was obtained as the input of scstGCN and other baselines. The results show that scstGCN outperforms all baselines on MBHD and HBCHD datasets from different species. It is worth emphasizing that scstGCN’s 25th percentile (0.7059 for MBHD and 0.6839 for HBCHD) of SSIM are even higher than the maximum of 75th percentile (0.6912 for MBHD and 0.6483 for HBCHD) of SSIM for state-of-the-art method iStar (Figure 4A). Certainly, the 50th percentile of scstGCN is also significantly lower than the minimum 25th percentile of iStar in terms of RMSE. In addition, the super-resolution gene expression predicted by iStar on the Visium HD datasets exhibit many extreme values, which may be linked to the number of genes (Figure 4A). As the number of genes increases, the gene expression data becomes sparser, making it more challenging to capture sufficient statistical information, which can lead to outliers for other baselines that don’t establish complex communication relationships between neighboring cells. This results in a more pronounced advantage for scstGCN in terms of the mean evaluation metrics, with a mean RMSE of 0.0462 and 0.0507 and SSIM of 0.7919 and 0.7692 for scstGCN across MBHD and HBCHD datasets, compared to RMSE of 0.0881 and 0.1267 and SSIM of 0.5735 and 0.4874 for iStar.

**Fig. 4.**
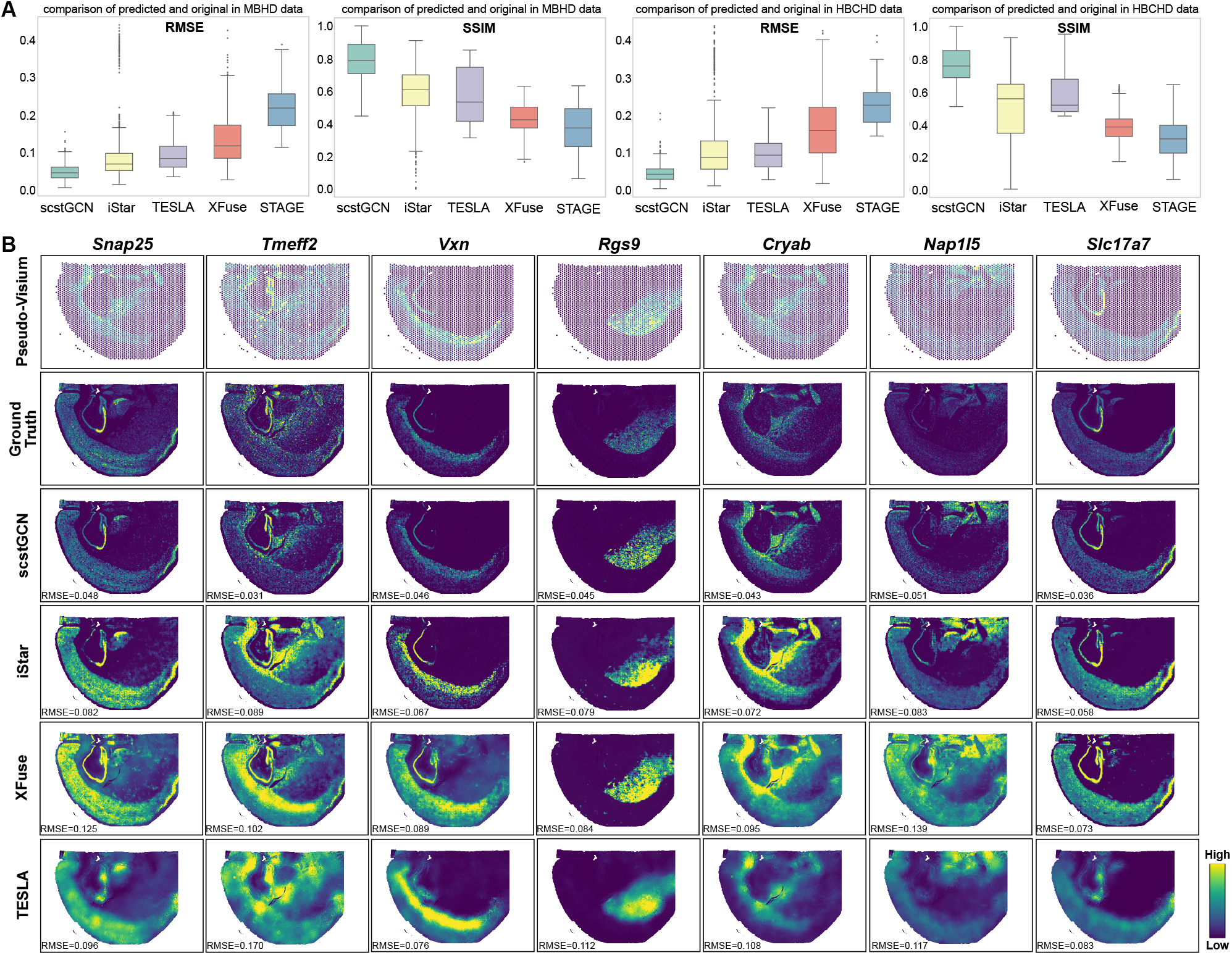
scstGCN enables accurately restore gene expression data for Mouse Brain (MBHD) and Human Breast Cancer (HBCHD) datasets from Visium HD technology. (**A**) Comparison of RMSE and SSIM between ground truth and predicted single-cell resolution gene expression by scstGCN, iStar, TESLA, XFuse, and STAGE on the MBHD and HBCHD datasets. (**B**) Spatial visualization of several genes having different spatial patterns for the ground truth and predicted data by scstGCN and other baselines in MBHD data.

In addition, due to the strong sense of pattern in the mouse brain, we spatially visualized several genes with different spatial patterns in ground truth and predicted profiles by scstGCN, iStar, XFuse, and TESLA in MBHD data (Figure 4B). Visually, the single-cell resolution gene expression profiles predicted by scstGCN is closer to the ground truth. iStar only performs a simple mapping from hierarchical histological features to gene expression, without considering the complex relationship between adjacent cells, which results in discontinuous predicted gene mapping and prone to blurred phenomenon. XFuse is misled by intense morphological similarities between different regions in histology image, resulting in poor prediction of low-expression regions of genes. TESLA comprehensively considers the physical distance between superpixels and the histological similarity, but does not deeply integrate gene expression with histological image, resulting in the overall smoothness of the predicted gene expression profiles, which deviates from the ground truth.

### scstGCN demonstrates superior overall performance on DLPFC datasets

Here, we evaluated the ability of scstGCN to predict super-resolution gene expression on the DLPFC tissue with 12 sections from 10X Visium platform. Since DLPFC data lacks super-resolution gene expression as labels, we conducted a post-processing step to precisely evaluate the performance of scstGCN and iStar numerically. Specifically, we aggregated the predicted super-resolution gene expression data by summing the gene expression values of superpixels covered by each 10X Visium spot, thus obtaining predicted gene expression at the spot-level. As we have conducted comprehensive comparisons using three Xenium datasets and two Visium HD datasets that have single-cell resolution gene expression as labels, and considering the long runtimes of XFuse and TESLA and the need for post-processing steps on the DLPFC data, so we only consider comparison with the state-of-the-art method iStar on the DLPFC data. We evaluated the prediction performance numerically by calculating the RMSE, Pearson correlation coefficient (PCC), and mean absolute error (MAE) between the original data and the predictions generated by scstGCN and iStar on the DLPFC dataset with 12 sections (Figure 5A and Supplementary Figure S3). The results indicate that scstGCN outperformed iStar across almost all genes in 12 sections with statistical significance (median: PCC = 0.83, RMSE = 0.08 and MAE = 0.06 for scstGCN; PCC = 0.67, RMSE = 0.17 and MAE = 0.09 for iStar).

**Fig. 5.**
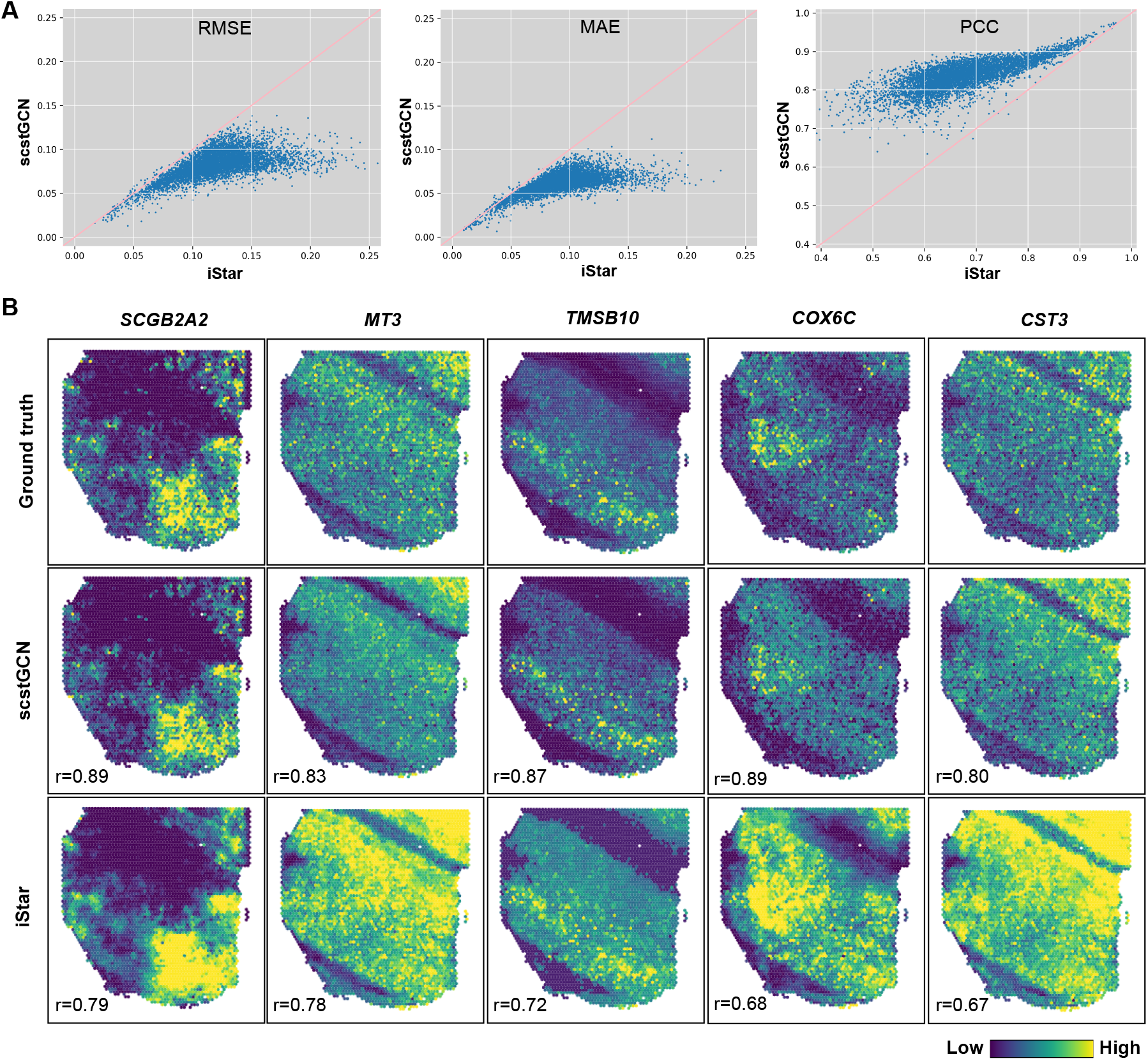
scstGCN demonstrates superior performance in the human dorsolateral prefrontal cortex (DLPFC) tissue compared to state-of-the-art method iStar. (**A**) Scatter plots of RMSE, MAE, and PCC between the original spot-level gene expression and “spot-level” gene expression obtained from the enhanced expression generated by scstGCN and iStar for the top 1000 highly variable genes selected across all sections in DLPFC tissue. Each dot in plots represents one of the 1000 genes. (**B**) Spatial visualization of several genes having different spatial patterns for the ground truth and predicted data by scstGCN and iStar in section 151507, respectively.

Similarly, we selected some highly variable genes that have different spatial patterns for spatial visualization to illustrate the advantages of scstGCN. The results show that the gene expressions predicted by scstGCN were closer to the ground truth, accurately recovering the original spatial patterns of the genes (Figure 5B). In contrast, the results from iStar visually restored the spatial patterns but showed high expression levels throughout the tissue compared to the ground truth. iStar exhibits excessive smoothing in local regions of gene expression, resulting in blurriness, while scstGCN provides more finer depiction of gene expression. iStar predicts gene expression by considering only the features of the superpixel itself, without leveraging the effective knowledge from neighboring superpixels. On the other hand, scstGCN enables the neural network to adaptively learn different gene expression levels corresponding to different regions by integrating spatial positional information into histological features. In addition, scstGCN can not only improve the resolution within the measured spot, but also predict single-cell resolution gene expression in non measured spot areas of histological images (Supplementary Figure S4).

To further demonstrate the superiority of scstGCN in maintaining the spatial structures, we performed clustering on the data obtained from scstGCN and iStar with spatial clustering methods AVGN, STAGATE, and GraphST, using Adjusted Rand Index (ARI) as the evaluation metric. The results show that the “spot-level” data obtained from scstGCN is better than iStar in all spatial clustering algorithms, ARI has a significant improvement (Supplementary Table S2). scstGCN retains the structure of the spatial domain tightly when enhancing the resolution of spatial gene expression.

### scstGCN can enhance the spatial patterns in the human DLPFC tissue

Next, we tested the ability of scstGCN to enhance the spatial patterns of genes in ST. We applied a tool called Sepal [39], which employs a diffusion-based model to identify transcripts exhibiting spatial patterns in the transcriptome, to DLPFC datasets before and after super-resolution prediction using scstGCN. Following the tutorial of Sepal, we ranked the original and super-resolution gene expression profiles in each section by the degree of spatial structure from distinct to random patterns respectively; then grouped top-ranked genes that had distinct spatial patterns into pattern families based on the similarity of their spatial organization. Finally, we conducted functional enrichment analysis for each section in DLPFC datasets using the Gene Ontology: Biological Processes (GO: BP) database [40]. Figure 6A shows a subset of transcript profiles in 151510 section that top-ranked in the super-resolution data but lower rankings in original data. The super-resolution expressions of these genes have distinct spatial structure. In contrast, the spatial structures are more diffuse in the original data. *TPT1* is pivotal in cell growth regulation and cancer progression, making it a promising target for therapies aimed at inhibiting tumor development and metastasis [41]. *COX6C* is vital for mitochondrial function, influencing cellular energy production and potentially impacting diseases like neurodegeneration and cancer [42]. *PTGDS* is crucial for inflammation and neuroprotection, making it a promising target for treating neurological and inflammatory conditions [43]. *RTN1* regulates neuronal morphology and function, impacting neurodegenerative disorders and neuronal regeneration [44]. The super-resolution data predicted by scstGCN helps researcher better discover spatial patterns of these genes, significantly improving their rankings.

**Fig. 6.**
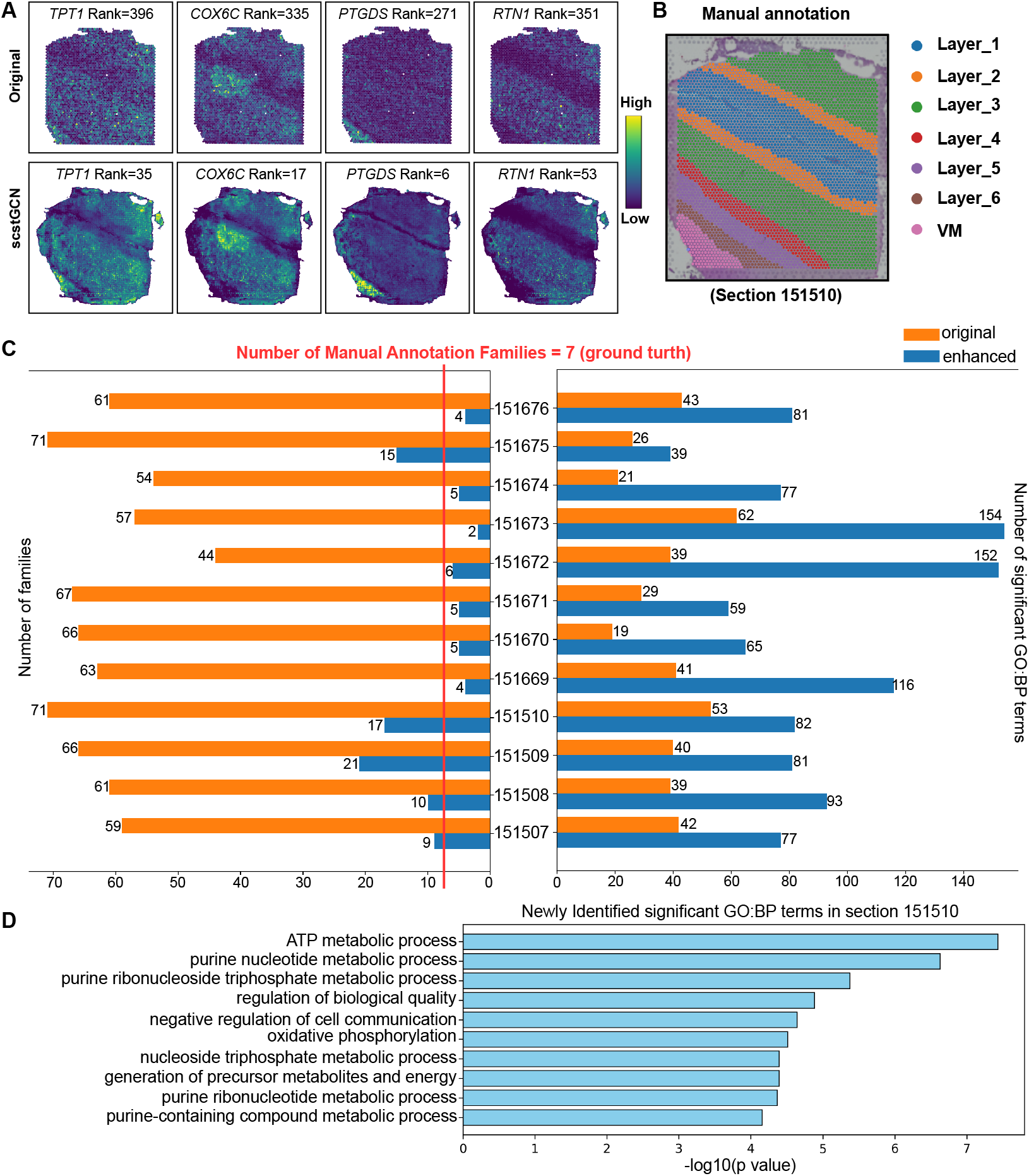
scstGCN can help find spatial patterns for DLPFC tissue and enrich more biologically significant pathways. (**A**) Some disease-related genes rank higher in super-resolution data predicted by scstGCN (below) because of distinct spatial patterns but rank lower in original Visium data (above). The head of each gene expression profile provides the names and specific ranking of the genes. (**B**) Manual annotation of hematoxylin-and-eosin-stained tissue image in section 151510. (**C**) Number of families divided according to similarity in spatial structure (left) and significant (*P*-value*<*0.05) GO:BP terms (right) for all sections. Orange represents the original Visium data, while blue represents the super-resolution data. (**D**) Top-10 significant GO:BP terms were only identified in the super-resolution data in section 151510.

Figure 6B shows that the DLPFC tissue has six cortical layers and white matter (WM) annotated manually. Next, we partitioned the top 100 ranked genes into pattern families using Sepal in both the original data and the super-resolution data across all sections (Figure 6C). In the original Visium data, the number of families partitioned by Sepal is much greater than the manually annotated count because low-resolution expression blurs the family patterns. The super-resolution gene expression data brings the number of families closer to manual annotations because it maximally restores the actual biological features of the tissue and enhances spatial patterns.

Finally, we conducted enrichment analysis in DLPFC tissue among the top 100 ranked genes. The results show that super-resolution data were enriched more GO: BP term across all sections (Figure 6C). Specifically, section 151672 enriched 113 new pathways, section 151673 enriched another 92 new pathways, and the least enriched section 151675 also identified 13 new pathways. Moreover, high-resolution DLPFC data revealed a variety of biological processes closely related to metabolic activities and regulatory networks (Figure 6D and Supplementary Figure S5), among which adenosine triphosphate (ATP) metabolic production, precursor and energy metabolites, and oxidative phosphorylation were closely related to schizophrenia [45, 46], which provides new perspectives and strategies for studying disease mechanisms and identifying therapeutic targets. By inferring super-resolution gene expression to accurately capture and analyze tissue structure details, scstGCN brings new advances to systems biology and biomedical research.

### Explore insights by super-resolution gene expression

We further analyzed the capability of scstGCN in super-resolution tissue structure segmentation and annotation. We applied scstGCN to the HER2ST data [29], which has a lower resolution than Visium. Specifically, we considered three consecutively cut sections from sample *H*, where manual annotations were only provided for section *H*_1. We used the spot-based ST data and histology image from section *H*_1 for training data, and then we used only the histology images from three sections as inputs to obtain the super-resolution gene expression. We used the output of the GCN as features for the k-means algorithm [47], and the resulting segmentation showed a strong concordance with the manual annotations (Figure 7A). scstGCN successfully separated In situ cancer, Breast glands, and Adipose tissue from manually annotated section *H*_1. Additionally, the tissue segmentation results have similar structure across the three sections, demonstrating that scstGCN has robust transfer learning capabilities and maintains consistency across different sections from the same sample.

**Fig. 7.**
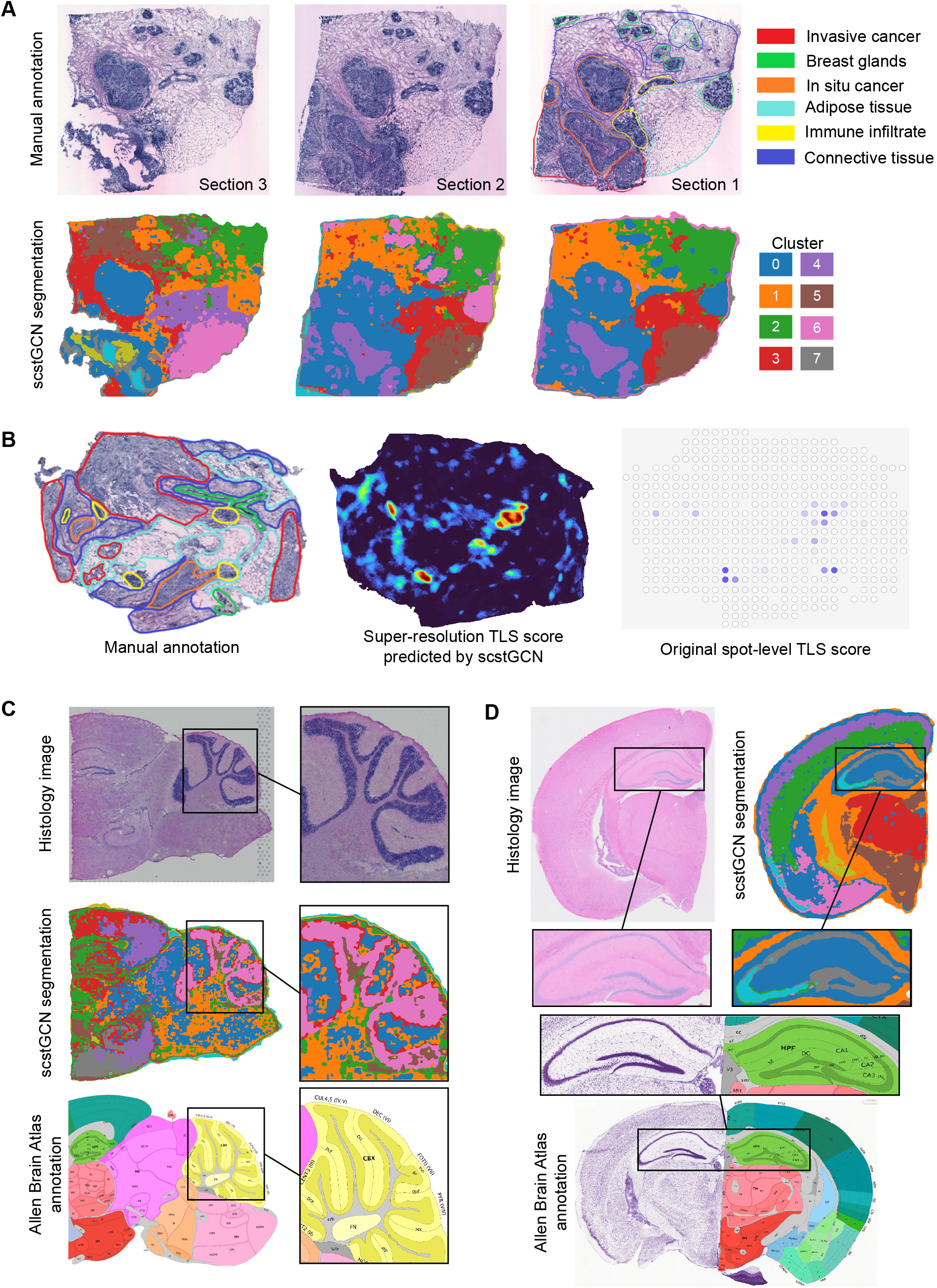
Exploring the capability of scstGCN for segmenting and annotating tissue structures at super-resolution across different types of datasets. (**A**) Annotating tissue architecture at superpixel-level using scstGCN on three consecutive tissue sections from sample *H* in the HER2ST breast cancer dataset. (**B**) Super-resolution gene expression obtained from section *G*_1 of HER2ST breast cancer dataset by scstGCN can facilitate the discovery of tertiary lymphoid structures (TLSs). (**C, D**) The tissue segmentation results based on super-resolution gene expression closely resemble the details of the Allen Brain Atlas. (**C**) mouse brain coronal cut data. (**D**) mouse brain sagittal cut posterior data.

Next, we explored the potential of scstGCN in detecting multicellular structures. Tertiary lymphoid structures (TLSs) refer to specific types of lymphoid tissue structures that form in non-lymphoid tissues [48]. Research indicates that TLSs may play a significant role in tumor immune surveillance and therapeutic responses. However, the resolution of ST data may not be sufficient to accurately capture TLSs because they are typically small and sparsely distributed lymphoid structures, especially under low signal-to-noise conditions. We calculated TLS scores separately on the super-resolution gene expression and Visium data of the section *G*_1 of another sample *G* in the HER2ST dataset by normalizing and averaging the TLS marker genes (Supplementary Table S3). The results show that TLSs detected from the Visium data are unusually sparse and have low density. On the contrary, super-resolution data predicted by scstGCN can detect more clearer and densely packed TLSs at a fine-grained characteristics (Figure 7B).

Finally, we applied scstGCN to MBS and MBC datasets to demonstrate that scstGCN is a generic tool not limited to a specific type of cancer or healthy tissue. We used the same method as in Figure 7A to perform tissue segmentation, and the results show that scstGCN could characterize fine-grained tissue structure with high-resolution (Figure 7C and D). The fine-grained tissue segmentation based on super-resolution gene expression predicted by scstGCN closely aligns with Allen Brain Atlas annotations. This further demonstrates the accuracy and reliability of scstGCN. When Allen Brain Atlas Annotation does not fully cover certain data, scstGCN serves as a new effective tool for brain research, opening new avenues for disease research and therapeutic strategies.

In addition, we also compared the ability of the spatial clustering algorithms STAGATE and GraphST to identify spatial domains with scstGCN, although both methods only identify spatial domains at the spot-level. The results show that STAGATE and GraphST did not show excellent spatial domain recognition performance on HER2ST data (Supplementary Figure S6). In contrast, scstGCN provides a better fine-grained super-resolution segmentation effect, which is consistent with the rough manual annotation. STAGATE and GraphST exhibited more pronounced effects in the specific regions identified in the MBC data, but the regions scstGCN identifies in MBS data may be more obvious (Supplementary Figure S7). scstGCN is not a tool specifically designed to identify spatial domains, which provides an idea for fine-grained high-resolution tissue segmentation.

Finally, we assessed the predictive performance of super-resolution gene expression in all of the aforementioned datasets. scstGCN outperformed iStar for virtually all genes across all datasets (Supplementary Figure S8).

### Assessing the impact of each module in scstGCN on predicted single-cell resolution gene expression results

In order to comprehend the reasons behind the performance improvement of scstGCN compared to other methods, we assessed the contribution of each module to scstGCN, including various feature map from mutimodal feature map (denoted as “UNI”, “LOC”, and “RGB” respectively) and the GCN module to scstGCN, as shown in Table 2. We systematically removed various combinations of four modules, including individual and combined removals. It is worth noting that the histological feature maps must retain at least one of the “UNI”, “LOC”, and “RGB” modules, and when the GCN is removed, we replace it with a feedforward neural network with four hidden layers, each containing 512 nodes, and ReLU as the activation function for the hidden layers was employed. From the numerical evaluation results of RMSE and SSIM in five Xenium datasets, keeping all modules intact provides the best alignment of the predicted super-resolution gene expression with the ground truth. Notably, removing the GCN module resulted in a significant performance drop across all datasets, only slightly better than iStar. Removing the UNI module led to an even more substantial decline.

**Table 2.**
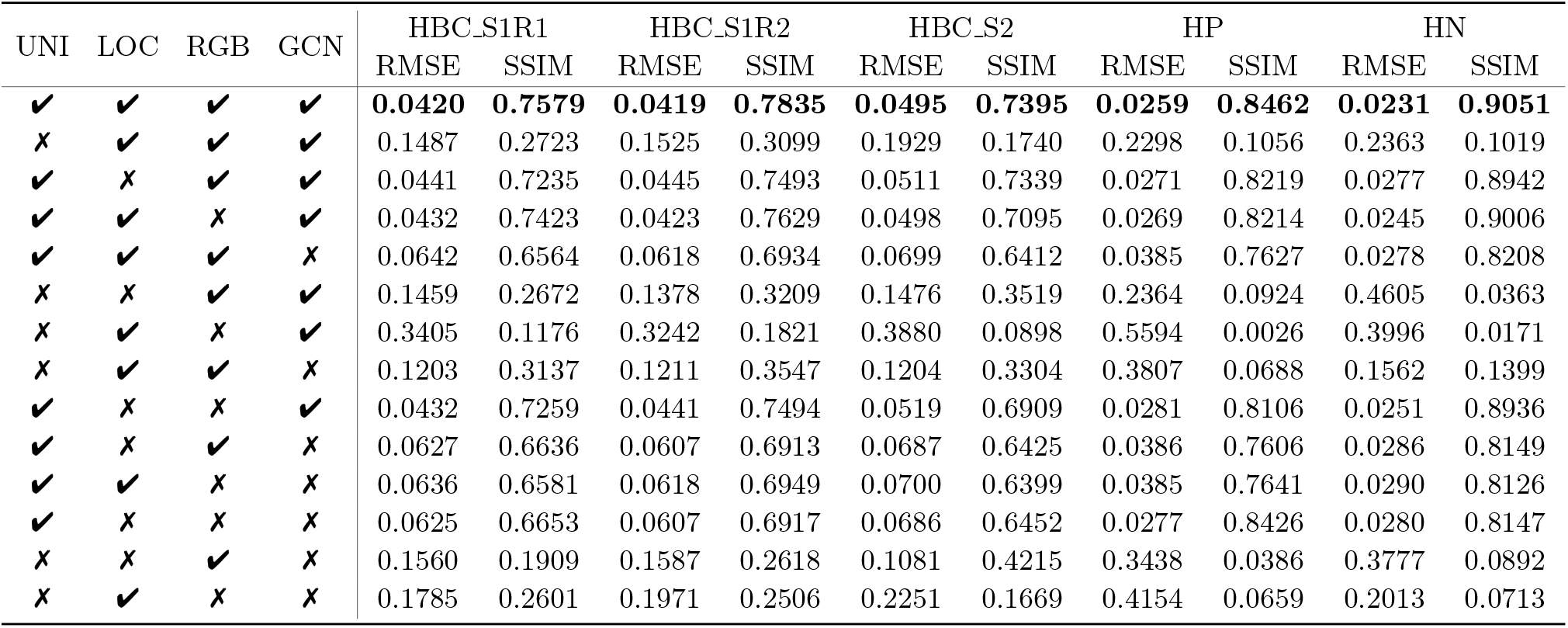
Ablation experiments on the HBC, HP, HN datasets from Xenium platform. The results showed the average RMSE and SSIM for scstGCN and its variants after removing specific modules in each dataset.

| UNI | LOC | RGB | GCN | HBC_S1R1 |  | HBC_S1R2 |  | HBC_S2 |  | HP |  | HN |  |
| --- | --- | --- | --- | --- | --- | --- | --- | --- | --- | --- | --- | --- | --- |
|  |  |  |  | RMSE | SSIM | RMSE | SSIM | RMSE | SSIM | RMSE | SSIM | RMSE | SSIM |
| ✓ | ✓ | ✓ | ✓ | **0.0420** | **0.7579** | **0.0419** | **0.7835** | **0.0495** | **0.7395** | **0.0259** | **0.8462** | **0.0231** | **0.9051** |
| ✗ | ✓ | ✓ | ✓ | 0.1487 | 0.2723 | 0.1525 | 0.3099 | 0.1929 | 0.1740 | 0.2298 | 0.1056 | 0.2363 | 0.1019 |
| ✓ | ✗ | ✓ | ✓ | 0.0441 | 0.7235 | 0.0445 | 0.7493 | 0.0511 | 0.7339 | 0.0271 | 0.8219 | 0.0277 | 0.8942 |
| ✓ | ✓ | ✗ | ✓ | 0.0432 | 0.7423 | 0.0423 | 0.7629 | 0.0498 | 0.7095 | 0.0269 | 0.8214 | 0.0245 | 0.9006 |
| ✓ | ✓ | ✓ | ✗ | 0.0642 | 0.6564 | 0.0618 | 0.6934 | 0.0699 | 0.6412 | 0.0385 | 0.7627 | 0.0278 | 0.8208 |
| ✗ | ✗ | ✓ | ✓ | 0.1459 | 0.2672 | 0.1378 | 0.3209 | 0.1476 | 0.3519 | 0.2364 | 0.0924 | 0.4605 | 0.0363 |
| ✗ | ✓ | ✗ | ✓ | 0.3405 | 0.1176 | 0.3242 | 0.1821 | 0.3880 | 0.0898 | 0.5594 | 0.0026 | 0.3996 | 0.0171 |
| ✗ | ✓ | ✓ | ✗ | 0.1203 | 0.3137 | 0.1211 | 0.3547 | 0.1204 | 0.3304 | 0.3807 | 0.0688 | 0.1562 | 0.1399 |
| ✓ | ✗ | ✗ | ✓ | 0.0432 | 0.7259 | 0.0441 | 0.7494 | 0.0519 | 0.6909 | 0.0281 | 0.8106 | 0.0251 | 0.8936 |
| ✓ | ✗ | ✓ | ✗ | 0.0627 | 0.6636 | 0.0607 | 0.6913 | 0.0687 | 0.6425 | 0.0386 | 0.7606 | 0.0286 | 0.8149 |
| ✓ | ✓ | ✗ | ✗ | 0.0636 | 0.6581 | 0.0618 | 0.6949 | 0.0700 | 0.6399 | 0.0385 | 0.7641 | 0.0290 | 0.8126 |
| ✓ | ✗ | ✗ | ✗ | 0.0625 | 0.6653 | 0.0607 | 0.6917 | 0.0686 | 0.6452 | 0.0277 | 0.8426 | 0.0280 | 0.8147 |
| ✗ | ✗ | ✓ | ✗ | 0.1560 | 0.1909 | 0.1587 | 0.2618 | 0.1081 | 0.4215 | 0.3438 | 0.0386 | 0.3777 | 0.0892 |
| ✗ | ✓ | ✗ | ✗ | 0.1785 | 0.2601 | 0.1971 | 0.2506 | 0.2251 | 0.1669 | 0.4154 | 0.0659 | 0.2013 | 0.0713 |

But one key point to emphasize is that removing UNI and relying solely on “LOC” and “RGB” modules for gene expression prediction will inevitably lead to a significant drop in performance. In contrast, if UNI is replaced with other pre-trained large models for extracting histological features, the differences will be minimal. To further demonstrate the contribution of our study, we compare scstGCN to state-of-the-art method iStar where both are using the same underlying ViT. We replaced the histological feature extraction of scstGCN with the HIPT hierarchical Transformer to make it consistent with iStar, and we refer to this method as “scstGCN+HIPT”. On the other hand, we also replaced iStar’s hierarchical histological feature extractor HIPT with UNI to make it consistent with scstGCN, which we refer to as “iStar+UNI”. We conducted a series of numerical evaluation experiments on the HBC datasets from Xenium platform and the MBHD and HBCHD datasets from Visium HD platform by comparing the prediction accuracy of those methods. The results indicate that replacing UNI with HIPT hierarchical Transformer did not significantly decrease the model’s performance compared to the original scstGCN. Even with the replacement of UNI with HIPT for feature extraction, our method (“scstGCN+HIPT”) still achieved higher median and mean values in RMSE and SSIM compared to iStar (Table 3 and Supplementary Figure S9). Specifically, compared to iStar on the mean, “scstGCN+HIPT” has increased by 42.5%, 55.0% on the RMSE,and 35.9%, 55.5% on the SSIM respectively on MBHD and HBCHD datasets, with similar improvements observed in three sections of HBC dataset (details in Table 3). In addition, the median comparison with iStar yielded results consistent with the mean. We also analyzed the difference in performance between iStar and “iStar+UNI”. For numerical evaluation metrics, “iStar+UNI” has a slight improvement in HBC S1R1 but even a slight decrease in HBC S1R1 and HBC R2 compared to iStar. To further verify the importance of GCN module, we set up a method to improve iStar, introducing GCN module to capture complex relationships between neighboring cells while keeping the upstream feature map acquisition part unchanged. The experimental results show that the introduction of GCN to capture complex relationships between neighboring cells significantly improves the prediction accuracy in iStar (Supplementary Figure S10 and Table S4).

**Table 3.**
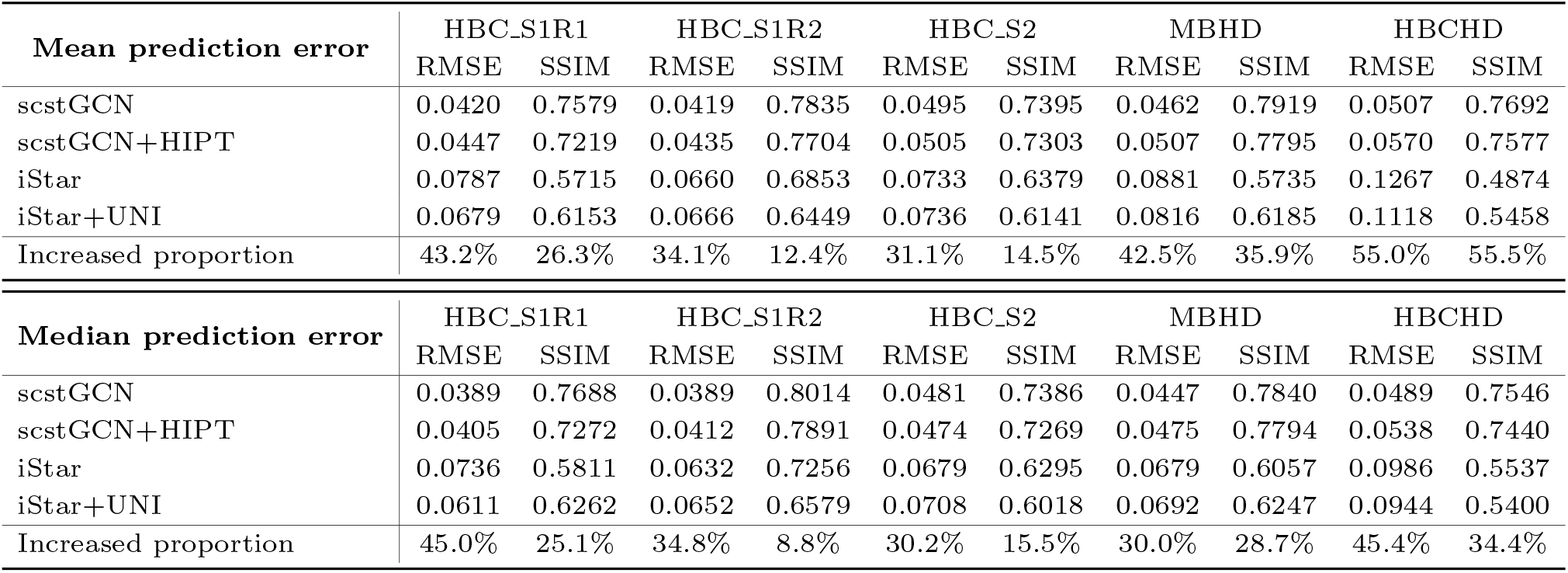
Comprehensively evaluate the performance of scstGCN and iStar, under the condition that both use the same pretrained ViT. **“Increased proportion”** represents the percentage of performance improvement of scstGCN+HIPT compared to iStar.

The experimental results indicate that there is a minimal performance difference in predicting single-cell resolution spatial gene expression between using UNI and other pre-trained large models as feature extractors for histology images. On the contrary, establishing complex communication relationships between neighboring cells using the GCN module is crucial for enhancing resolution of gene expression. Relying on histological features to predict gene expression is the cornerstone, and the GCN module of establishing complex communication relationships between neighboring cells plays a crucial role in improving accuracy. Additionally, removing the “LOC” and “RGB” modules from the histological feature maps also reduced performance to varying degrees.

## Conclusion

Spatial transcriptomics is a revolutionary technology with tremendous potential for elucidating cellular composition, understanding molecular interactions between tissue components, and advancing disease research [49]. Despite its advantages, the low resolution or inability to cover the entire transcriptome limits their application. Computational predictions of high-resolution gene expression are effective solution, but existing methods often fail to accurately capture the full complexity of cellular features. Here, we developed scstGCN, a GCN-based method that leverages a weakly supervised learning framework to integrate multimodal information and then infer super-resolution gene expression at single-cell level. scstGCN can accurately enhance gene expression from the spot-level to the superpixel-level, and it can predict expression both outside the spots and in external tissue sections.

However, the datasets used in this study are limited to a narrow range of sequencing platform types. In future studies, we consider introducing diverse data types from multiple sequencing platforms. Our ideal solution is to develop a unified framework designed to enhance the spatial resolution of gene expression across various data types derived from different sequencing technologies. In addition, scstGCN currently lacks the capability to integrate multi-omics data. In future studies, we aim to incorporate single-cell RNA sequencing (scRNA-seq) data and effectively fuse it with ST data and histological images using an attention mechanism. By combining multi-omics data at the informative feature representation level, we expect to enhance the accuracy and robustness of scstGCN’s single-cell resolution gene expression predictions.

Identifying genes that display spatial expression patterns is a crucial step in characterizing the complex tissue landscapes in spatial transcriptomics studies. Due to low resolution and high noise levels, original transcriptomics data makes it challenging to identify biologically significant genes with spatial patterns. We demonstrated that scstGCN can help identify spatial patterns of disease-related genes, which is of significant importance for understanding disease mechanisms by analyzing all sections from DLPFC tissue. Enrichment analysis in DLPFC tissue has shown that scstGCN allows for the enrichment of more biologically significant pathways, thereby providing deeper insights into biological processes. The ability of scstGCN to annotate tissues at super-resolution demonstrates great potential, showing consistency with manual annotations while providing higher granularity.

In summary, scstGCN provides a robust approach for ST data enhancement, including super-resolution gene expression generation, identification of spatial patterns of genes, enrichment of biologically significant pathways, and tissue structures segmentation. In addition, we quantified the runtime and memory consumption of scstGCN. scstGCN is the second fastest method, only slightly higher than iStar. And scstGCN consumes the least Video Random Access Memory (VRAM) (Supplementary Table S5).

## Acknowledgments

The work was supported in part by the National Natural Science Foundation of China (62262069 and 12101026), and the Yunnan Talent Development Program - Youth Talent Project.

## Notes

### Competing Interest Statement

The authors have declared no competing interest.

